# Synergistic effects of the next generation insecticide flupyradifurone with a fungal pathogen

**DOI:** 10.1101/2023.07.31.551355

**Authors:** Daniel Schläppi, Adam Al-Hashemi, Vaneeza Wasif, Florent Masson, Nathalie Stroeymeyt

## Abstract

The global insect decline is largely driven by agricultural pesticides, whose detrimental effects are further amplified by synergistic interactions with other stressors. Whilst there is increasing evidence of adverse impacts of the next-generation insecticide flupyradifurone (FPF), initially marketed as safe for pollinators, its effects on non-pollinators and potential interactions with other stressors remain poorly understood. Here we investigate the impacts of chronic exposure to FPF alone and in combination with the entomopathogenic fungus *Metarhizium brunneum* in a non-target arthropod: the ant *Lasius niger*. Our results show that FPF concentrations upwards of 100 ppm increased worker mortality. Furthermore, while sublethal (≤ 50 ppm) FPF exposure alone did not affect survival, it increased susceptibility to fungal infection, indicating a synergy between the two stressors. This new report of a synergistic interaction between FPF and pathogens raises further concerns about the impact of novel pesticides and could have important implications for insect conservation.

## Introduction

Insects are important indicators for ecosystem health and provide vital ecosystem services such as pollination, pest control, and nutrient cycling^1,2^. However, global insect biodiversity and abundance are being lost at alarming rates^3,4^. Recent research identified agrochemicals such as neonicotinoid insecticides as a major driver of these declines^5,6^. The detrimental impacts of neonicotinoids on non-target organisms^7-10^ led to restrictions in the use of clothianidin, thiamethoxam, and imidacloprid in the European Union. Following the neonicotinoid ban, new pesticides were introduced to the market, including the butenolide insecticide flupyradifurone (FPF). Although structurally different, neonicotinoids and FPF target the same receptor and both disrupt the central nervous system of insects, making them highly effective against a broad range of pests^11,12^. Upon introduction, FPF was advertised as “bee safe” because of its ecotoxicological profile. However, recent evidence has uncovered that FPF does pose a risk to pollinators, as exposure to environmentally relevant doses of FPF leads to detrimental sublethal effects, including impaired cognitive abilities, motion coordination deficits, hyperactivity and apathy, as well as impacts on flight and foraging behaviours^13-17^. Such sublethal impacts frequently elude detection in risk assessments which are typically based on acute toxicity assays.

A major concern associated with the use of agrochemicals are their interactions with other stressors, as synergistic interactions may amplify their environmental significance. Traditional agrochemicals have been demonstrated to interact synergistically with a variety of stressors such as other pesticides, pathogens, malnutrition, or climate change, leading to increased host mortality^18-22^. By contrast, there have been conflicting reports regarding interactions between FPF and other pesticides^23-25^, and so far, there is no evidence for an interaction between FPF and disease^26,27^. Furthermore, the great majority of studies examining the direct and indirect impacts of FPF have focused on bees, whilst other non-target insects such as ants have been overlooked, in spite of their ubiquity and the essential ecosystem services they provide^28^. Here we aimed to address these gaps by (i) investigating the direct effects of FPF exposure on ant survival and (ii) testing for potential indirect effects via interaction with disease susceptibility. To do so, we first determined the susceptibility of the black garden ant *Lasius niger* to exposure to a range of FPF concentrations. We then used exposure to field-realistic, sublethal FPF doses in combination with the fungal pathogen *Metarhizium brunneum* to investigate potential interactive effects between the two stressors.

## Methods

*L. niger* colonies were initiated using newly mated queens collected in Berlin (July 2021). The colonies were raised at 25°C and 65% humidity, with a 12h day/night cycle, *ad libitum* supply of water and honey water (15% mass fraction of honey) and weekly provision of *Drosophila hydei*.

### Flupyradifurone susceptibility test

We sampled 120 workers from each of four stock colonies and randomly split them into eight subsets of fifteen workers (N=480 ants across 32 subsets). Each subset was kept in a separate Petri-dish (⌀ = 50 mm) with fluon-coated walls and sealed with parafilm. The floor of each Petri-dish was covered with a thin layer of plaster of Paris to which 1 mL of water was added to maintain humidity. After one day of acclimatisation, each colony’s subsets were pseudo-randomly allocated to the control treatment or one of seven FPF concentration treatments. Although field-realistic exposure doses remain to be determined for ants^9,29^, data provided by the Environmental Protection Agency (EPA) showed residues up to 36 ppm in flowers and up to 4.3 ppm in nectar^30^. Hence, we considered 5 ppm to be a conservative estimate for field-realistic dose and designed our experimental concentration range around that value. Fresh FPF feeding solutions were prepared on each feeding day by mixing different proportions of a 30% honey water solution, a 1000 ppm FPF stock solution and MilliQ water to produce 8 different solutions with final concentrations of 15% honey water each and 0 (control), 0.5, 1, 5, 10, 50, 100 or 500 ppm FPF (Sigma-Aldrich PESTANAL® analytical standard, 99.5% purity). At the start of the experiment and every 3 days thereafter, each subset received a fresh FPF feeding solution provided in a 200 μL Eppendorf tube closed with cotton wool. Survival was checked daily for two weeks by counting and removing dead ants. All survival checks were conducted blind to treatment.

### Flupyradifurone and fungus synergy test

To test for potential interactive effects between FPF and pathogens, ants were chronically exposed to sublethal FPF doses for ten days, and then subsequently challenged with a fungal pathogen. We used the generalist entomopathogenic fungus *Metarhizium brunneum* (strain MA275, KVL 03-143). *M. brunneum* is common in soil environments and naturally infects *L. niger*, which display a variety of physiological and behavioural defences against it^31-33^. A preliminary test was conducted to confirm that FPF does not directly interfere with fungal germination (supplementary Fig. 1).

We sampled 132 workers from each of five stock colonies and distributed them evenly across 3 separate Petri-dishes, resulting in a total of 15 Petri-dishes containing 44 ants each (total ants N = 660). The three Petri-dishes from each colony were pseudo-randomly assigned to one of three FPF treatment groups (0, 5 and 50 ppm of FPF, that is, control, low, and high sublethal doses as determined in experiment 1). Feeding solutions were freshly prepared and provided every 3 days as described above. After ten days, 40 workers from each Petri dish were randomly selected and split into two groups of 20, which were randomly allocated to two pathogen treatments (fungus-exposed and sham-exposed). This resulted in a full factorial design involving six treatment groups (three FPF treatments ×two pathogen treatments) with 5 replicates each (one per stock colony). Accordingly, the experiment was organised into 5 successive experimental blocks, each involving a single stock colony and one replicate for every treatment. In each block, all sham- and fungal exposures were performed in a single session lasting less than two hours. The 20 ants of each Petri-dish were exposed consecutively, while the order in which the Petri-dishes for all treatments were exposed was varied pseudo-randomly across all blocks to mitigate temporal effects.

Fungal and sham exposures were performed using standard procedures^33^: *M. brunneum* was cultured on Sabouraud-Dextrose Agar (SDA) plates at 24°C until sporulation. Conidiospores were harvested in 0.05% Triton-X100, washed twice and adjusted to 10^9^ spores/mL in 0.05% Triton-X100 to create a spore stock suspension. The stock was diluted by half with 0.05% Triton-X100 resulting in a fungal suspension with a final spore concentration of 5×10^8^ corresponding to the median lethal dose LD50 (unpublished data). Ants were placed on ice, then exposed to 0.3 μL of the treatment solution (fungal suspension or sham solution of 0.05% Triton X-100 only) pipetted onto their gaster. They were then left to dry on filter paper for 1 minute and subsequently transferred individually into individual Petri-dishes (⌀ = 35 mm, N = 600) with fluon coated walls and a water tube (200 μL Eppendorf tubes closed with cotton wool). These Petri-dishes were then sealed with parafilm and placed in the incubator for two weeks with daily survival checks conducted blind to treatment.

### Statistical Analyses

All statistical analyses were performed in using R v4.2.2^34^. Survival analyses were performed using Cox proportional hazards mixed-effect models using R packages *survival*^35^ and *coxme*^36^. For the FPF susceptibility test treatment group was used as a categorical fixed effect and Petri-dish and stock colony as random effects. For the FPF and fungus synergy test, fungal exposure group, FPF concentration and their one-way interaction were used as categorical fixed effects and Petri-dish, stock colony and exposure block as random effects. Fixed effect significance was tested using two-tailed Wald chi-square tests. *Post-hoc* contrasts, defined using package *multcomp*^37^, were used for pairwise comparisons of experimental groups. P-values were corrected for multiple comparisons using the Benjamini-Hochberg (BH) method.

## Results

### Flupyradifurone susceptibility test

FPF concentration had a significant effect on ant mortality (Fig. 1; Cox proportional hazards mixed effect model, effect of FPF concentration: χ^2^ = 31.67, df = 7, p < 0.001). *Post-hoc* pairwise comparisons revealed that concentrations of 100 ppm and above led to a significant increase in ant mortality compared to the control (100 ppm vs. 0 ppm: z = 2.81, p=0.017; 500 ppm vs. 0 ppm: z = 4.97, p < 0.001). By contrast, concentrations of 50 ppm and below did not increase mortality compared to the control (z < 2.08; p > 0.09 in all pairwise comparisons); these were used as sublethal concentrations in experiment 2. The absence of lethal effects in concentrations of 50 ppm and below cannot be attributed to the ants restricting their food intake to limit their exposure to the pesticide, as feeding volumes were only found to decrease at the highest FPF concentration tested (500 ppm, see supplementary Fig. 2).

**Figure 1.**
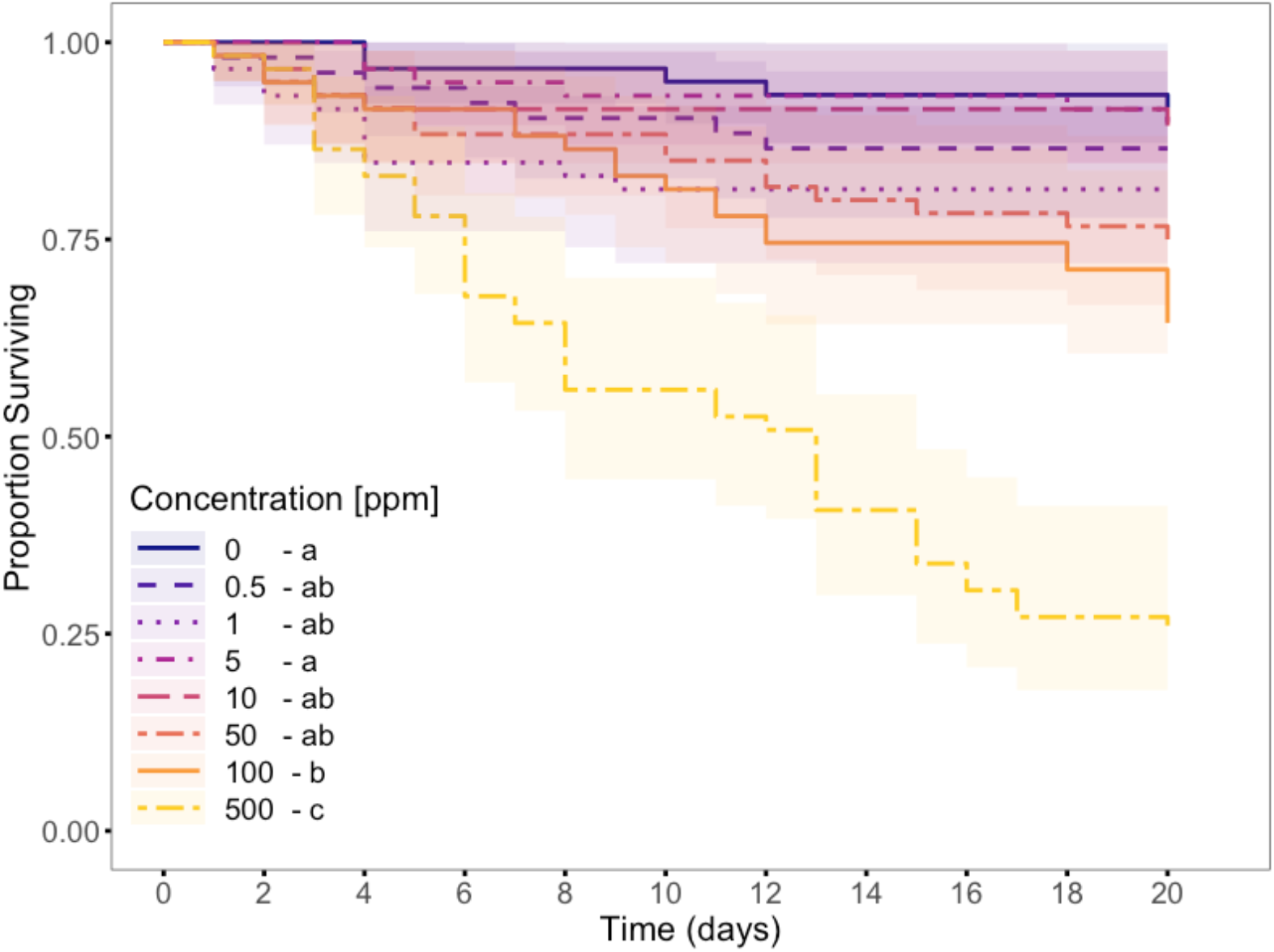
Flupyradifurone susceptibility. Lines represent the proportion of *L. niger* workers surviving as a function of time during chronical exposure to different concentrations of FPF (total sample size: N=467). Shaded areas represent the 95% confidence interval around each line. Concentration had a significant effect on survival (Cox proportional hazards model; χ^2^ = 31.699, df = 7, p < 0.001). Results of pairwise comparisons are indicated by letters in the legend (same letter: p > 0.05; different letter: p ≤ 0.05 in *post hoc* tests with BH correction).

### Flupyradifurone and fungus synergy test

Challenge with the pathogenic fungus *M. brunneum* increased ant mortality in all three FPF exposure conditions (Fig. 2a; Cox proportional hazards mixed effect model, effect of fungal exposure: χ^2^ = 57.49, df = 1, p < 0.001; pairwise comparisons of fungus-treated vs. sham-treated ants with BH correction, no FPF: Hazard Ratio (HR) = 1.69, z = 2.36, p = 0.028; FPF 5 ppm: HR = 3.56, z = 5.04, p < 0.001; FPF 50 ppm: HR = 3.96, z = 5.94, p < 0.001). Furthermore, although FPF alone did not affect ant survival (χ^2^ = 3.84, df = 2, p = 0.15), there was a significant interaction between FPF treatment and fungal exposure (χ^2^ = 8.29, df = 2, p = 0.016). More specifically, prior sublethal exposure to FPF significantly increased the fungus-induced mortality compared to the control (Fig. 2b; pairwise comparisons of Hazard Ratios with BH correction, control vs. FPF 5ppm: z = 2.22, p = 0.032; control vs. FPF 50ppm: z = 2.65, p = 0.016; FPF 5ppm vs. 50ppm: z = 0.31, p = 0.76). This indicates that infections with *M. brunneum* are even more likely to be lethal if the ants have previously been chronically exposed to a sublethal FPF dose (Fig. 2b).

**Figure 2.**
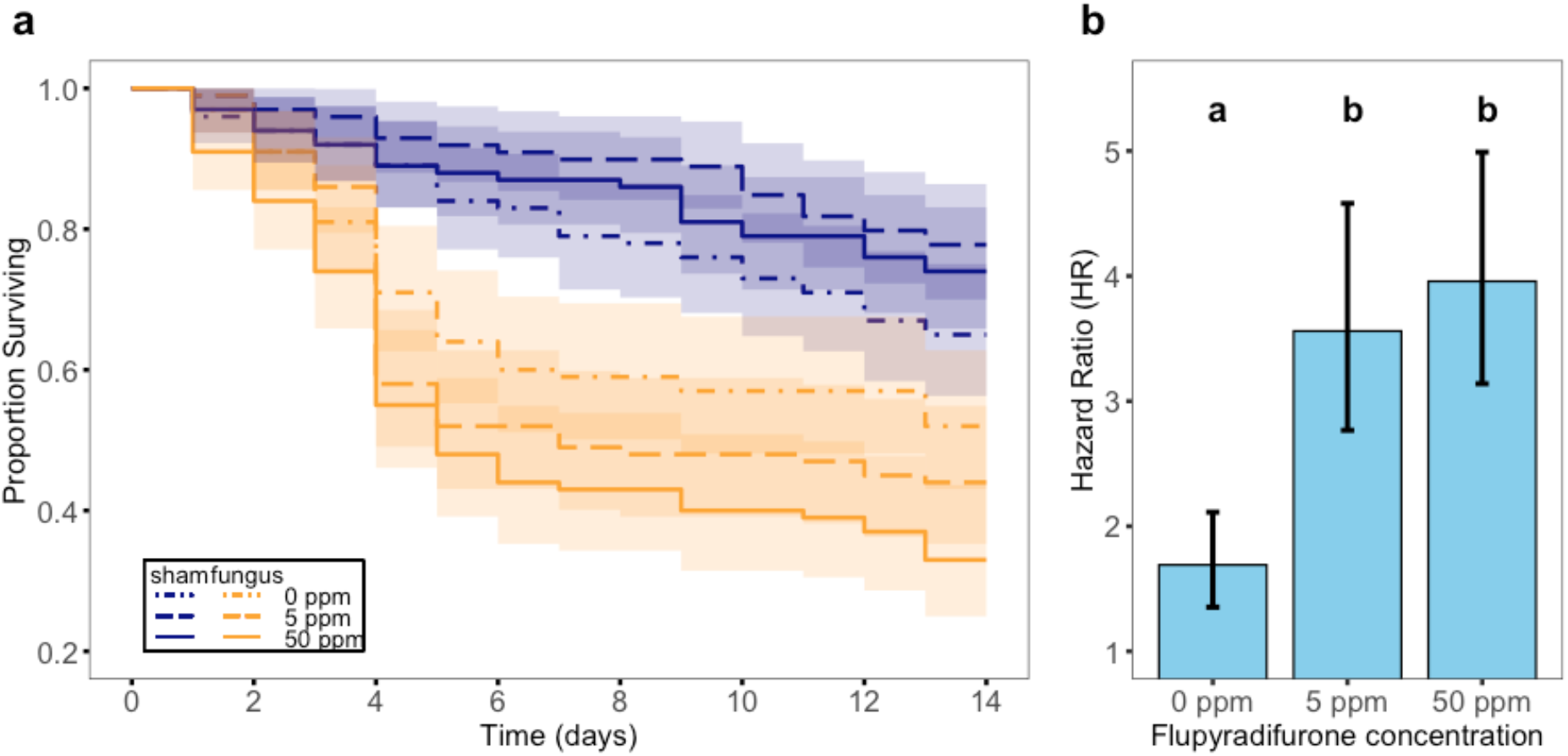
Interactive effects between Flupyradifurone and *M. brunneum*. a)Proportion of *L. niger* workers surviving as a function of time following exposure to *M. brunneum* spores (yellow lines) or to a sham solution (blue lines) after ten days chronic exposure to different concentrations of flupyradifurone (0, 5, and 50 ppm; dashdotted, dashed and solid lines, respectively). Shaded areas represent the 95% confidence interval around each curve. b) Hazard Ratio of fungus-treated ants relative to sham-treated ants in the three FPF exposure conditions. Bars and whiskers represent model estimates and standard errors, respectively. Cox proportional hazards mixed effect model, interaction FPF exposure X fungal exposure: χ^2^ = 8.22, df = 2, p = 0.016. Results of pairwise comparisons are indicated by letters (same letter: p > 0.05; different letter: p ≤ 0.05; *post hoc* contrast with BH correction).

## Discussion

This study assessed the direct effect of chronic FPF exposure on the survival of black garden ants (*L. niger*), revealing increased mortality at concentrations of 100 ppm and above. Furthermore, we demonstrated an indirect effect of FPF on ants at lower, sublethal concentrations (≤ 50 ppm), as the challenge with *M. brunneum* was more likely to be lethal following chronic exposure to sublethal FPF doses, indicating a synergistic interaction between the two stressors. Overall, the results of this study raise further concerns about the long-term impacts of novel pesticides on the health of insects.

While chronic exposure to FPF concentrations upwards of 100 ppm resulted in increased worker mortality, concentrations of 50 ppm or lower showed comparable mortality rates to the control group and were thus classified as sublethal. However, the direct impact of FPF on ants in the field remains unclear as the exact exposure of ants to FPF in their natural environment has yet to be quantified. Residues in pollen and nectar are considered to be the main exposure routes for bees and their reported residue values are as high as 68 ppm for pollen and 21.8 ppm for extrafloral nectaries^38^. However, studies of honey bees typically use 4 ppm as field-realistic doses because nectar collected by returning foragers contained 4.3 ppm of FPF^16,30^. Our sublethal concentrations for ants (0.5 -50 ppm) cover a comparable range, suggesting that FPF residue values in the field may not increase ant mortality via direct toxicity. Nonetheless, future studies should quantify true field-realistic exposure for ant colonies over the course of a year, based on a combination of all relevant acute and chronic exposure routes, which includes contact with spray droplets, contaminated soils, foliage, and water, as well as foodborne exposure^9,29,39^.

At sublethal concentrations, FPF exposure alone did not affect ant survival, but it increased mortality when combined with *M. brunneum* exposure, indicating that FPF may induce indirect sublethal effects in non-target hosts at field-realistic concentrations via a synergistic interaction with pathogens. This contrasts with previous studies in bees, which did not detect FPF-induced changes in survival rates to fungal or viral pathogens^26,27,40^. However, our findings are consistent with findings from traditional neonicotinoids in ants, as imidacloprid was shown to increase the susceptibility of ants to infection by the fungus *Beauveria bassiana*^41^.

The synergistic effect between FPF and *M. brunneum* suggests that FPF may affect the behavioural or biochemical immune response of the ants. However, as we did not record the ants’ behaviour or measure their immune function, we cannot identify the underlying mechanism for this interaction. On one hand, FPF could cause a direct immune suppression or trigger a resource allocation trade-off between immune functions and detoxification^42^, as has been shown in honey bees exposed to neonicotinoids^18,43,44^. On the other hand, as FPF disrupts the function of the central nervous system, it could also interfere with individual and collective behaviours that mitigate the risk of fungal infection in ants, such as self-grooming, allo-grooming or chemical disinfection^31^. Future studies involving entire colonies will be required to determine whether FPF impairs colony-wide social immunity mechanisms that protect colonies against epidemics and how individual level effects impact the colony as a whole^45-47^.

Overall, this study contributes to the understanding of stressor interactions by providing evidence for a synergistic effect between FPF and pathogens in a non-target organism, and it highlights the ecotoxicological risk posed by novel insecticides. Our findings are particularly concerning as they suggest that FPF may pose a comparable threat to beneficial insects as banned neonicotinoids, despite its initial perception of relative safety for bees. This stresses the importance of considering sublethal, long-term effects as well as interactive effects between multiple stressors when assessing the risks of agrochemicals, rather than only evaluating direct mortality at low doses. Furthermore, our findings revealed a synergistic interaction between FPF and pathogens in ants, which contrasts with the absence of such an effect in previous studies on honey bees. This shows that the use of bees as surrogates for other non-target organisms may be inadequate when evaluating the risk of agrochemicals, and suggests that other economically and ecologically important arthropods such as ants should be included alongside bees as representative model organisms.

## Limitations of the study

This study demonstrates the synergistic effects of FPF and fungal pathogens in a controlled laboratory setting, which may not account for the full range of environmental factors that ants encounter in natural settings. Additionally, the study focuses on *L. niger* ants and one exemplary fungal model pathogen, hence generalizations to other species should be made with caution. The absence of behavioural and immunological assessments leaves the underlying mechanisms behind the observed synergy speculative and further research is required to explore these aspects.

## Acknowledgements

We would like to thank Prof Nicolai Vitt Meyling from University of Copenhagen for providing *M. brunneum*. Financial support was provided to DS by the Swiss National Science Foundation (P2BEP3_195575 & P500PB_206883) and to N.S. by the European Research Council (ERC Starting Grant ‘DISEASE’, no. 802628).

## Author contributions

D.S. conceived the study and designed it with N.S. and F.M; D.S., V.W., and A. AH. conducted the research and performed the experiments with inputs from N.S. & F.M.; N.S. provided laboratory space and materials; D.S., V.W., and A. AH. analysed the data with inputs from N.S.; D.S. wrote the manuscript based on a first draft by A. AH, with contributions from all authors. All authors edited and approved the manuscript.

## Data accessibility statement

The supplementary files, raw data and code supporting the results are archived on figshare and can be accessed via DOI: 10.6084/m9.figshare.25782330

## Supplemental information titles and legends

Document S1. containing details on the antifungal assay including supplementary Fig. 1 and details on the quantification of food uptake and supplementary Fig. 2.

## Supplementary Materials

### Antifungal assay: Metarhizium brunneum growth inhibition test

Flupyradifurone acts by specifically impairing the nervous system of insects (Nauen et al., 2015) and thus, no antifungal activity is expected from this agrochemical. Nonetheless, any results of combined exposure treatments would be difficult to interpret or even invalidated if flupyradifurone did inhibit the growth of *M. brunneum*. To test the ability of flupyradifurone to inhibit *M. brunneum* germination we used a disk-diffusion assay as described in Balouiri et al. (2016). Briefly, SDA plates were inoculated with 50 μL of 10^9^/mL *M. brunneum* conidiospore suspension. Then, filter paper disks (6 mm, Cytiva Life Sciences) were soaked with 15 μL of flupyradifurone solutions (10, 100, 1000 ppm), distilled water (0 ppm; negative control) or 5% sodium hypochlorite (positive control) and put on the agar using forceps. Plates were sealed, incubated for 48h at 24°C and photographed to measure the growth inhibition diameter. Even at the highest flupyradifurone dose no grow inhibition zone was visible, suggesting that *M. brunneum* germinated normally. Our findings are in line with findings for neonicotinoids, which have been used successfully in studies with fungal pathogens where no negative effects on conidia germination, conidia production and vegetative growth of Metarhizium fungi were detected (Cramer, 2020; Neves et al., 2001, Santos et al., 2007).

**Supplementary Figure 1.**
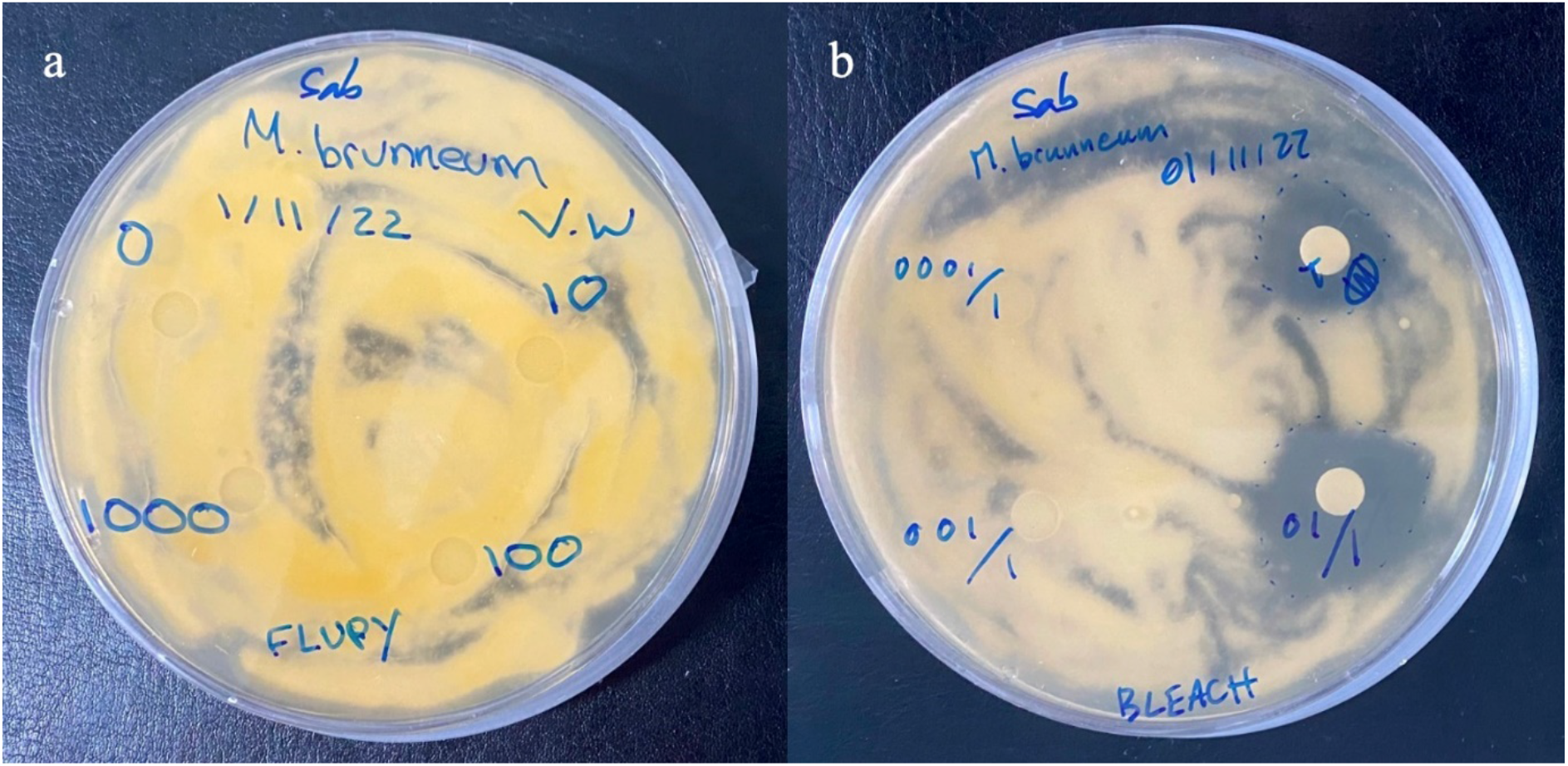
Antifungal assay. Disk diffusion method to test for germination inhibition of (a) flupyradifurone at 0, 10, 100 and 1000 ppm and (b) sodium hypochlorite (positive control).

### Quantification of food uptake

#### Methods

Pesticide contamination can modulate feeding motivation and food uptake, as shown for honey bees (Kessler et al., 2015; Siviter & Muth, 2022; Wu et al., 2021). Thus, to determine whether disparities in survival may be due to differences in food consumption, we quantified honey water uptake over a day at different FPF concentrations. For this purpose, we quantified fluorescence of homogenized ants that had access to a feeding solution containing fluorescein dye (Jové et al., 2020). The feeding solutions were prepared as described in the main manuscript, but with a fraction of water replaced by fluorescein sodium salt (Sigma-Aldrich) stock solution at 2% in MiliQ water. Four treatment solutions with final concentrations of 15% honey water, 0.01% fluorescein and varying concentrations of FPF (0, 5, 50 or 500 ppm, that is control, low, mid, and high treatment) were prepared. Workers were sampled from 5 source colonies and split into 4 subsets per colony in a stratified random way. Subsets were kept in petri dishes in the incubator for three days without food. Subsequently, each subset was assigned pseudo-randomly one of the four treatments groups. Final sample size was 106, 108, 105 and 102 workers in the control, low, mid and high treatment respectively. Workers had access to the solution for 24h before being freeze killed at -80°C for 30 min. Samples were then homogenized individually in racked collection tubes (Qiagen) containing 100 μL of phosphate-buffered saline (PBS) buffer and a 2 mm glass bead each. Homogenization was performed using a tissue lyser at 30 Hz for two runs of 1 minute with a plate rotation between the two runs. Following 2 minutes of centrifugation at 3000 rpm, 20 μL of the supernatant and 80 μL PBS buffer were transferred into 96-well plates. Fluorescence was measured on a SpectraMax iD5 plate reader with excitation at 485 nm and emission at 535 nm. Each measure was averaged from 4 reading points 1.5 mm apart, with 400 ms integration time and a gain of 500 volt.

Quantification was done using a standard curve, which was obtained using an aliquot of the control treatment feeding solution containing 0.01% fluorescein and 0 FPF, which was treated identically to the meals that were served to the ants (same light and temperature conditions throughout the duration of the experiment, and subsequent freezing). Reference standard curves were prepared from the retained aliquot by making a serial dilution for 8 solutions containing 5, 2.5, 1.25, 0.625, 0.3125, 0.15625, 0.078125, or 0 μL of 0.01% fluorescein treatment solution mixed with PBS to a final total volume of 100 μL. An unfed ant treated identically to the samples was used as a blank. To calculate the meal volumes, mean fluorescence value of blanks was subtracted from all other samples and consumed volume was extrapolated from the standard curves.

Statistical analyses were performed in R. A generalized linear mixed model (GLMM) with a negative binomial distribution was used to test for differences in meal size depending on FPF concentration. Treatment was included as fixed effect and petri dishes, colonies as well a block as random effects. Post hoc testing p-values were adjusted using the Benjamini-Hochberg method to correct for multiple comparisons.

#### Results

The estimated consumed volume of honey water over 24 hours, was significantly different among treatments (χ^2^ = 7.853, df = 3, p = 0.049; supplementary figure 2). Pairwise comparisons using Tukey adjustment indicated that ants in the treatment with the highest FPF concentration (500 ppm) consumed lower volumes (0.06 ± 0.07 μL) compared to ants in the low treatment (0.17 ± 0.14 μL; p = 0.048; fig 2). The remaining pairwise comparisons with the control and the mid treatment (0.159 ± 0.14 μL and 0.15 ± 0.14 μL respectively) were not significant.

**Supplementary Figure 2.**
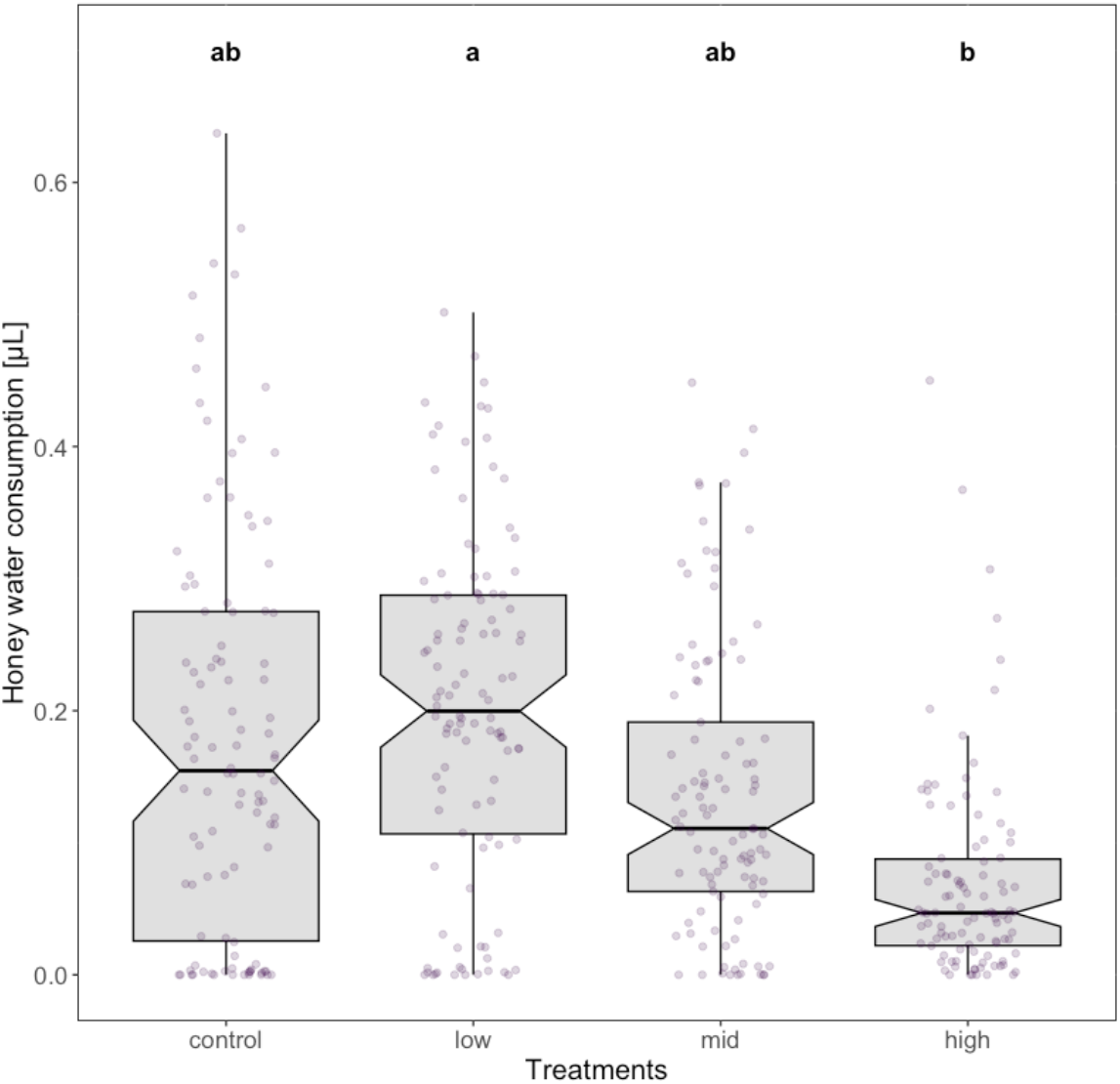
Quantification of food uptake. Volume (μL) of honey water consumed by *Lasius niger* workers in 24h at different concentrations of FPF (control = 0, low = 5, mid = 50, high = 500 ppm). Boxplots are shown with the inter-quartile-ranges (box), medians (black line in box) and outliers (dots). Transparent dots represent individual data points and bold letters (a,b) indicate significant differences (*p* < 0.05) between treatments (Tukey post hoc test).

In contrast to honey bees, which respond to field-realistic concentrations of FPF by a decreased food intake and erratic foraging behaviour (Hesselbach et al., 2020; Wu et al., 2021), we found that ants do not alter their food uptake if FPF is present in food. Even at 500 ppm the food uptake was comparable to the controls. Consequently, our results suggest that ants are not repelled by sublethal concentrations of FPF, i.e. below 50 ppm. The absence of a repellent effect implies that the ants are prone to collect contaminated food from the environment and sharing it among nestmates, with potentially severe consequences for the colony. The insecticide is likely to reach all members of the colony, including the queen. Even though the queen might have some protection via the colony or potentially superior detoxification compared to workers (Schläppi et al., 2020), she will still get exposed repeatedly over extended periods and thus might face a trade-off between detoxification and reproduction (Schwenke et al., 2016).

